# To what extent are the terminal stages of sepsis, septic shock, SIRS, and multiple organ dysfunction syndrome actually driven by a prion/amyloid form of fibrin?

**DOI:** 10.1101/057851

**Authors:** Douglas B. Kell, Etheresia Pretorius

## Abstract

A well-established development of increasing disease severity leads from sepsis through septic shock, SIRS, multiple organ dysfunction syndrome and cellular and organismal death. We argue that a chief culprit is the LPS-induced anomalous coagulation of fibrinogen to produce a form of fibrin that is at once inflammatory, resistant to fibrinolysis, and underpins the disseminated intravascular coagulation commonly observed in sepsis. In particular, we argue that the form of fibrin produced is anomalous because much of its normal α-helical content is transformed to β-sheets, as occurs in established amyloidogenic and prion diseases. We hypothesise that these processes play a major role in the passage along the above pathways to organismal death, and that inhibiting them would be of great therapeutic value, a claim for which there is emerging evidence.

## Introduction

Sepsis is a disease with high mortality [1–4]. However, the original notion of sepsis as the invasion of blood and tissues by pathogenic microorganisms has long come to be replaced, in the antibiotic era, by the recognition that in many cases the main causes of death are not so much from the replication of the pathogen *per se* but from the host’s ‘innate immune’ <u>response</u> to the pathogen (e.g. [5–8]). In particular, microbial replication is not even necessary (and anyway most bacteria in nature are dormant [9–13]), as this response is driven by very potent [14] inflammagens such as the lipopolysaccharides (LPS) of Gram-negative bacteria and equivalent cell wall materials such as lipoteichoic acids from Gram-positives [15]. To this end, such release may even be worsened by antibiotic therapy [16–19]. In unfavourable cases, this leads to an established cascade (Fig 1) [20] in which the innate immune response, involving pro-inflammatory cytokines such as IL-6, IL-8, MCP-1, TNF-α and IL-1β [21], becomes a ‘cytokine storm’ [22–26] leading to a ‘systemic inflammatory response syndrome’ (SIRS) [27–32], septic shock [3], apoptotic [33] and necrotic [34] cell death, multiple organ failure [35] (MOF, also known as Multiple Organ Dysfunction Syndrome, MODS [36; 37]), and finally organismal death. All of the above is well known, and may be taken as a non-controversial background.

**Figure 1:**
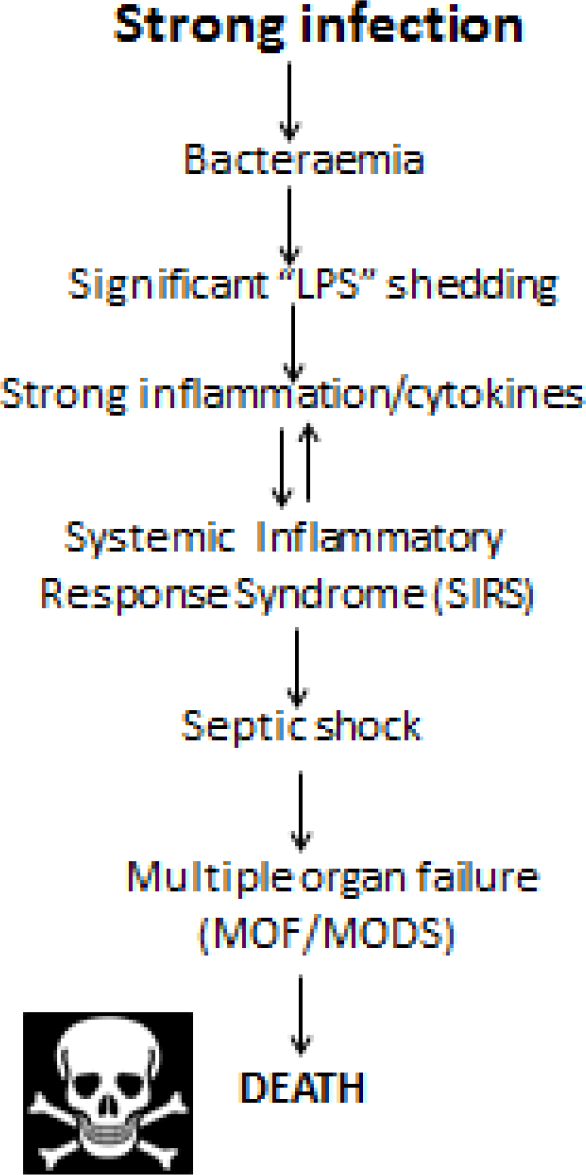
A standard cascade, illustrating the progression of infection through sepsis, SIRS and death.

Most recently [38], definitions of sepsis have come to be based on organ function and the Sequential (Sepsis-Related) Organ Failure Assessment (SOFA) Scores [39]. These latter take into account the multisystem nature of sepsis, and include respiratory, coagulatory (but only based on platelet counts), hepatic, cardiovascular, renal and CNS measurements. A SOFA score of 2 or greater typically means at least 10% mortality. Specifically, sepsis is defined as a life-threatening organ dysfunction caused by a dysregulated host response to infection. Septic shock is defined as a subset of sepsis in which underlying circulatory and cellular metabolism abnormalities are profound enough to increase mortality substantially.

**Table 1:**
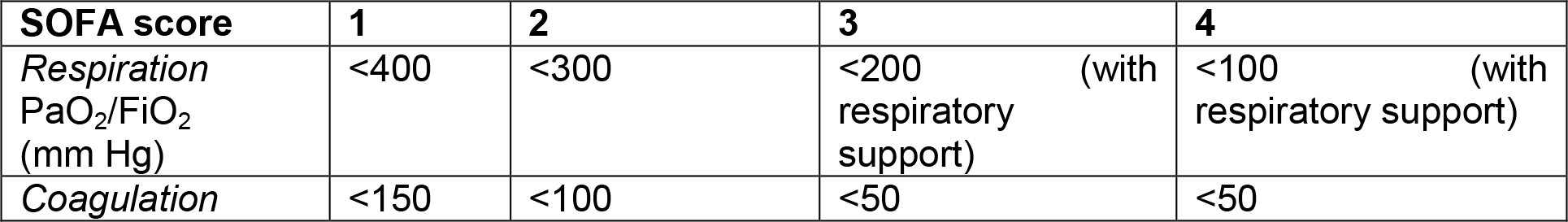
(based on [39]) shows the potential values that contribute to the SOFA score. *catecholamine and adrenergic agents administered for at least 1h; doses in μg.(kg.min)^−1^.

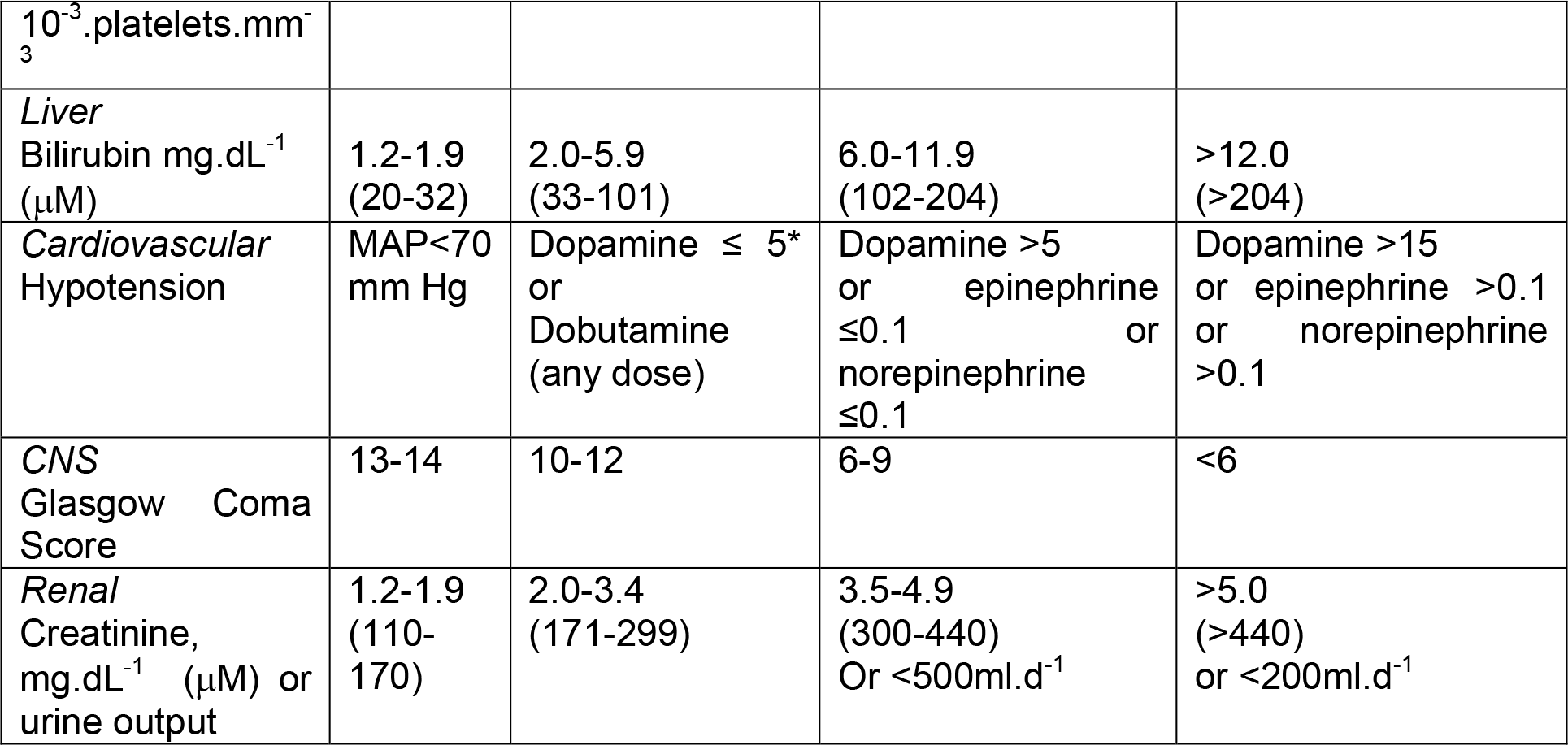

Absent from Fig 1 and the usual commentaries of this type are any significant role of coagulopathies, although these too are a well-established accompaniment of SIRS/sepsis [40–49], and they are our focus here. They form part of an emerging systems biology story (e.g. [13; 50–57]) in which iron dysregulation and an initially minor infection is seen to underpin the aetiology of many chronic inflammatory diseases normally considered (as once were gastric ulcers [58]) to lack a microbial component.

Here we develop the somewhat different case <u>for those conditions that are recognised</u> as involving a genuine initial microbial invasion and sepsis and inflammation driven (in particular) by the cell wall components of bacteria, although we note that the same kinds of arguments apply to viruses [59] and to other infections.

## Normal blood coagulation and coagulopathies

There are two main pathways of activation of ‘normal’ blood coagulation to form a clot, as occurs e.g. in response to wound healing. They have been expertly reviewed many times (e.g. [60–65], are known as ‘intrinsic’ and ‘extrinsic’, and are diagrammed in Fig 2. In short, after damage to e.g. blood vessel, collagen is exposed and **factor VII** leaves the circulation that comes into contact with tissue factor (TF), forming a complex called <u>***TF-FVIIa***</u> (complex shown in blue (block A) in Fig 2). This complex activates both factors IX and X (shown in orange). **Factor VII** is also activated by various molecules, including **thrombin**, **factor XIa**, **factor XII** and **factor Xa** (green arrows in Fig 2). Factor Xa and its co-factor Va form the prothrombinase complex (shown in purple (block B) in Fig 2) and activates thrombin via prothrombin (block C in Fig 2). Finally, the terminal stages of the coagulation pathway happens, where a cross-linked fibrin polymer is formed as a result of fibrinogen (typically present in plasma at 2-4 g.L^−1^) conversion to fibrin and cross-linking due to the activation of factor XIII, a transglutaminase. Thrombin activates factor XIII into factor XIIIa which finally acts on fibrin to form the cross-linking between fibrin molecules to form an insoluble fibrin clot. This fibrin clot, when viewed under a scanning electron microscope, consists of individually visible fibrin fibres, discussed in the next paragraphs (see Fig 3A for a representative healthy clot structure, created when thrombin is added to plasma (e.g. [54; 66–68]).

**Figure 2.**
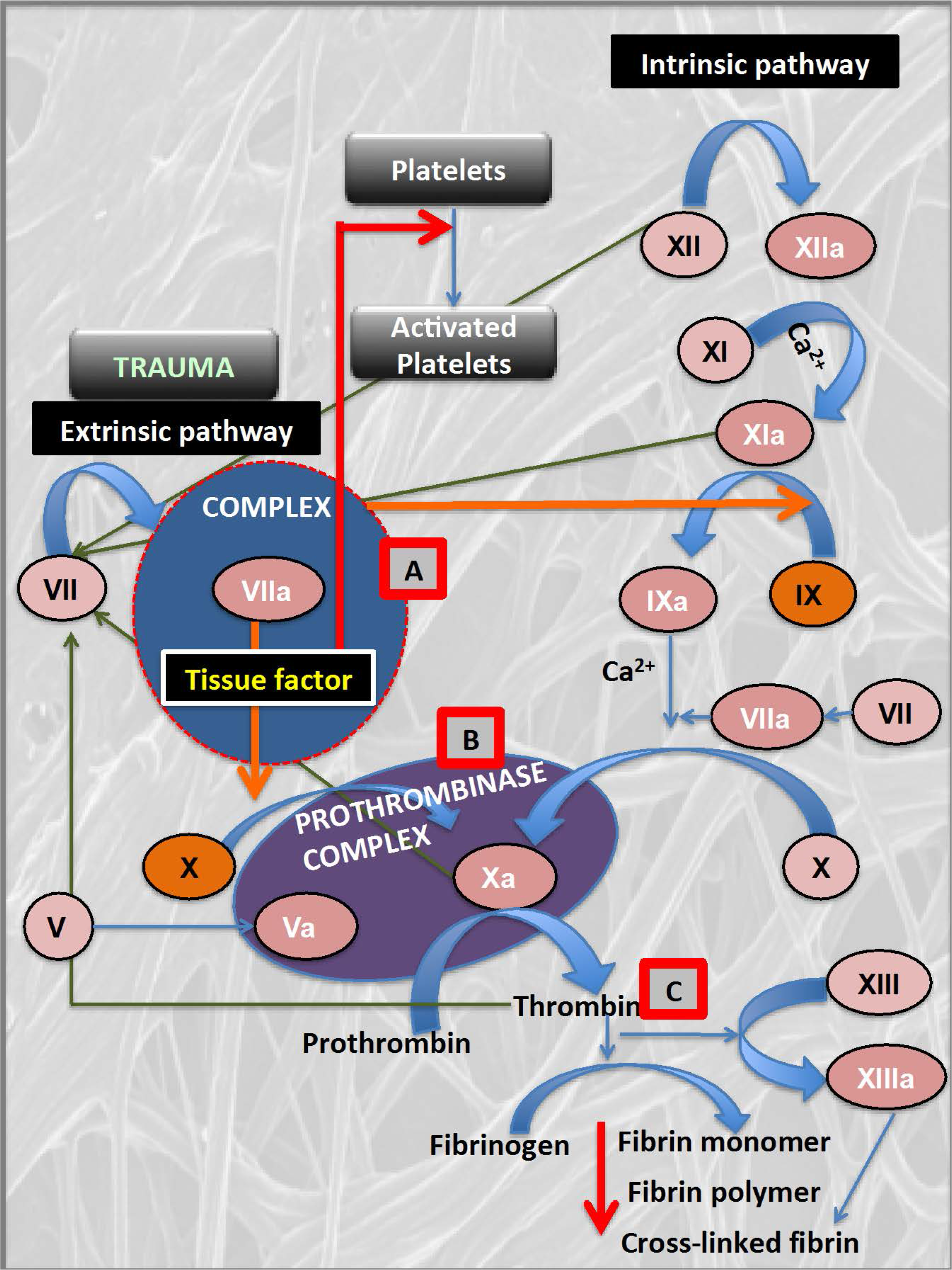
The coagulation pathway, with specific reference to facture VII and the tissue factor and factor VIIA complex, TF-FXIIa (block A) the prothrombinase complex (block B) and the activation and action of thrombin (block C).

The normal picture of fibrinogen polymerisation involves the removal of two fibrinopeptides from fibrinogen, which is normally rich in helices, leading to its self-association via knobs and stalks, but with otherwise no major changes in secondary structure.

Coagulopathies occur when the rate of clot formation or dissolution is unusually fast or slow, and in the case of chronic inflammatory diseases these seem largely to coexist as <u>hyper</u>coagulation and <u>hypo</u>fibrinolysis, arguably implying a common cause [54]. In a series of papers, we have shown in a number of diseases such as stroke [69–71], type 2 diabetes [68; 72], Alzheimer’s [73–75], and hereditary haemochromatosis [67], that instead of their usual ‘spaghetti-like’ appearance the fibrin clots induced by added thrombin adopted the form of ‘dense matted deposits’. The same kinds of effect could also be induced by unliganded iron [67; 76–78], although no molecular explanation was (or could be) given.

## Endotoxin-induced ‘disseminated intravascular coagulation’

Endotoxin (LPS) may also induce a runaway form of hypercoagulation (e.g. [41; 79–93]) known as disseminated intravascular coagulation (DIC). There is significant evidence, now that DIC is reasonably well defined [37; 94–96], that it can lead directly to multiple organ failure and death ([97], and see below). We hypothesise here that the form of clotting in DIC in fact involves autocatalytic β-amyloid formation (which is consistent with the faster clot formation in the presence of endotoxin [73]), and that this in particular is a major contributor to the various stages of sepsis, SIRS, MODS and ultimately organismal death.

## Prions. protein free energies, and amyloid proteins

Although it was originally shown that at least some proteins, when denatured and renatured, could revert to their original conformation [98; 99], implying that this was (isoenergetic with) the one of lowest free energy, this is now known not to be universal. Leaving aside chaperones and the like, one field in which proteins of the same sequence are well known to adopt radically different conformations, with a much more extensive β-sheet component (that is indeed thermodynamically more stable) is that of prion biology [100; 101]. The PrP^c^ and PrP^sc^ conformations, and the catalysis of the conversion to itself by the latter of the former are very well known. The key point for us here, however, is indeed that this definitely implies (e.g. [100–107]) that proteins that may initially fold into a certain, ostensibly ‘native’, conformation can in fact adopt stable and more β-rich conformations of a lower free energy, separated from that of the original conformation by a potentially significant energy barrier.

## Amyloid-like conformational transitions in fibrin(ogen)

As mentioned, the general view is that no major secondary structural changes occur during
normal fibrin formation. However, we know of at least three circumstances in which fibrin <u>can</u> (i.e. is known to) adopt a β-rich conformation. These are (i) in the case of specific mutant sequences of the fibrinogen a chain [108–114], (ii) when fibrin is stretched mechanically beyond a certain limit [115–121], (iii) when formed in the presence of certain small molecules, including bacterial lipopolysaccharide (LPS) [57; 122; 123]. Thus it is well established that fibrin can form β-rich amyloids, although it is assumed that conventional blood clotting involves only a ‘knobs and stalks’ mechanism without any major changes in secondary structure (e.g. [60–65; 124; 125]. We hypothesise here that the ‘dense matted deposits’ seen earlier are in fact β-rich amyloids, and that it is this coagulopathy in particular that contributes significantly to the procession of sepsis along or through the cascade of toxicity outlined in Fig 1.

In particular, thioflavin T (ThT) is a stain whose fluorescence (when excited at 440-450nm or so) is massively enhanced upon binding to β-rich amyloids (e.g. [126–135]). Fig 3 A, B show SEM pictures of ‘normal’ clots and dense matted deposits, respectively, while Fig 3C,D show how ThT stains clotted platelet-poor plasma from a patient with an inflammatory disease (type 2 diabetes) much more strongly than the equivalent plasma from a healthy control.

**Figire 3:**
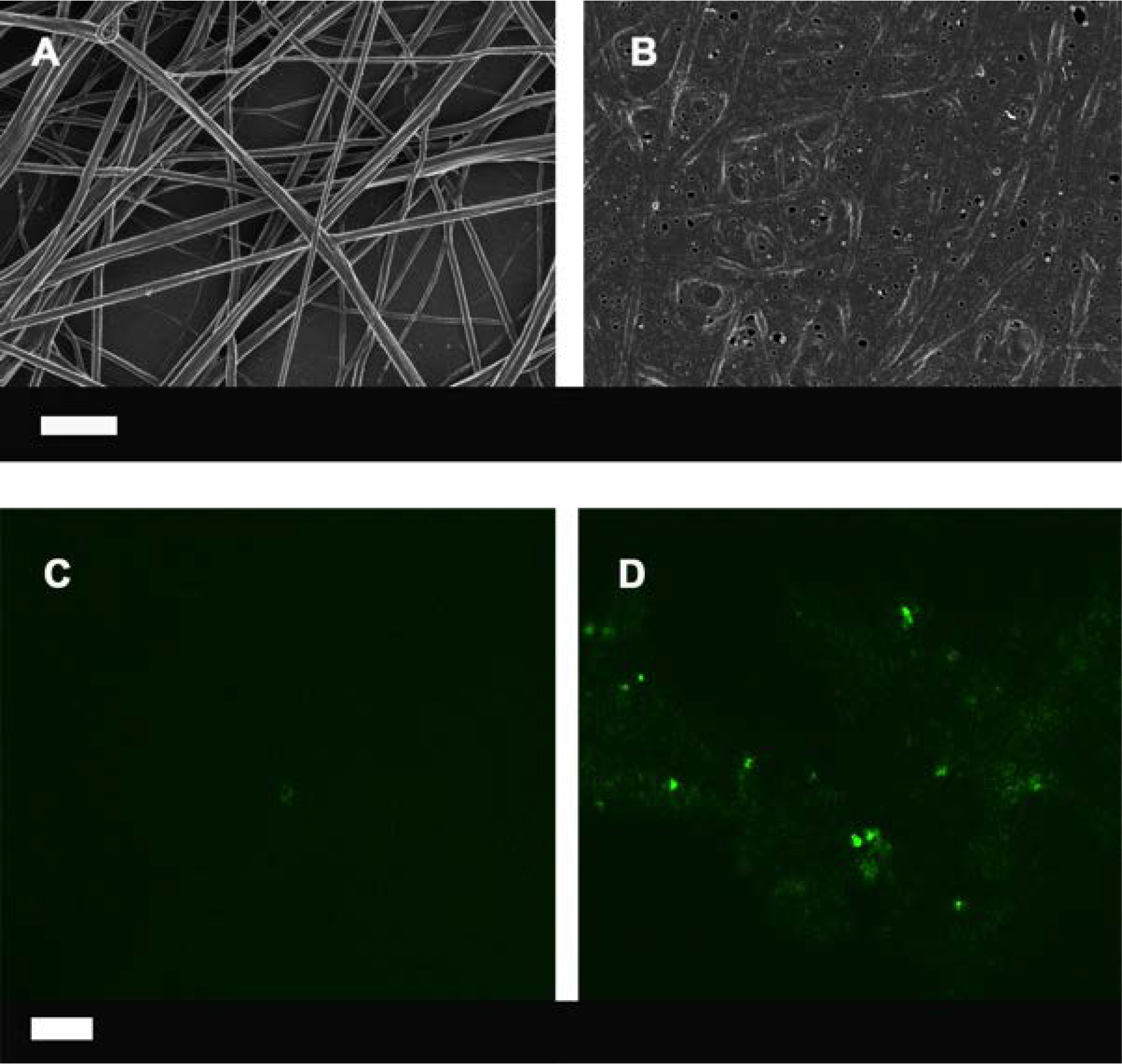
The results of thrombin-mediated blood clotting: **(A)** Normal fibrin fibres, **(B)** the same but taken from a patient with type II diabetes **(C)** ThT confocal ith no LPS, **(D)** ThT confocal with LPS. Scale bars of **A** and **B**: 1 μm and **C** and **D** 10 μm

## Inflammatory nature of fibrin

The fact that fibrin itself is or can be inflammatory is well established (e.g. [84; 136–141]), and does not need further elaboration; our main point here is that in none of these studies to date has it been established whether (or to what extent) the fibrin is in a β-amyloid form or not.

## Further evidence for the ‘trigger’ role of LPS in large-scale amyloid formation

In our previous studies [122], we found that LPS (endotoxin) at a concentration of just 0.2ng.L^−1^ could trigger the conversion of some 10^8^ times more fibrinogen molecules [122], and that the fibrin fibres so formed were β-amyloid in nature. (A very large amplification of structural molecular transitions could also be induced by LPS in a nematic liquid crystal [142–144].) Only some kind of autocatalytic processes can easily explain this kind of polymerisation, just as occurs in prions.

## Cytotoxicity of β-amyloids

This is so well known (e.g. [73; 145–153]) as barely to need rehearsing, although the relative toxicities of soluble material, protofibrils, fibrils and so on is less well understood [154], in part because they can equilibrate with each other even if added as a ‘pure’ component (of a given narrow MW range).

## Sequelae consistent with the role of p-amyloids in the ‘sepsis cascade’ to organ failure and death

If vascular amyloidogenesis really is a significant contributor to the worsening patient conditions as septic shock moves towards MOF/MODS and death, with the cytotoxic p-amyloids in effect being largely responsible for the multiple organ failure, then one might expect it to be visible as amyloid deposits in organs such as the kidney (whether as biopsies or post mortem). It is certainly possible to find evidence for this [155–160], and our proposal is that such amyloid should be sought using thioflavin T staining in autopsy tissue.

## How might this understanding lead to improved treatment options?

Over the years there have been many high-profile failures of therapies for various aspects of
severe inflammation, sepsis, septic shock, and SIRS. These include therapies aimed at endotoxin itself (Centoxin) [161–163], and the use of recombinant activated protein C [164], and of Drotrecognin alfa [163; 165–167]. Anti-cytokine and anti-inflammatory treatments have also had (at best) mixed results [168; 169].

However, the overall picture that we have come to is given in fig 4. This implies that we might hope to stop the progress of the sepsis/SIRS/MODS cascade at any (preferably several [170]) of a number of <u>other</u> places, including via iron chelation [50; 51; 171; 172], the use of anti-inflammatories, of anticoagulants such as heparin [140], and of stimulants of fibrinolysis [173]. The success of heparin [174; 175] (see also [176–178]) is especially noteworthy in the context of the present hypothesis, though it may have multiple (not simply directly anti-coagulant) actions [179; 180]. It is also noteworthy that HDL cholesterol is a protective against sepsis [55; 181–183] (HDL are anti-oxidant [184] and anti-inflammatory [185] and can also mop up endotoxin [186–188]), so the beneficial role of certain statins in sepsis [189; 190] should be seen in the context of their much more potent anti-inflammatory role [50] than any role involved in lowering overall serum cholesterol. Phospholipid emulsions may also serve [191].

**Figure 4:**
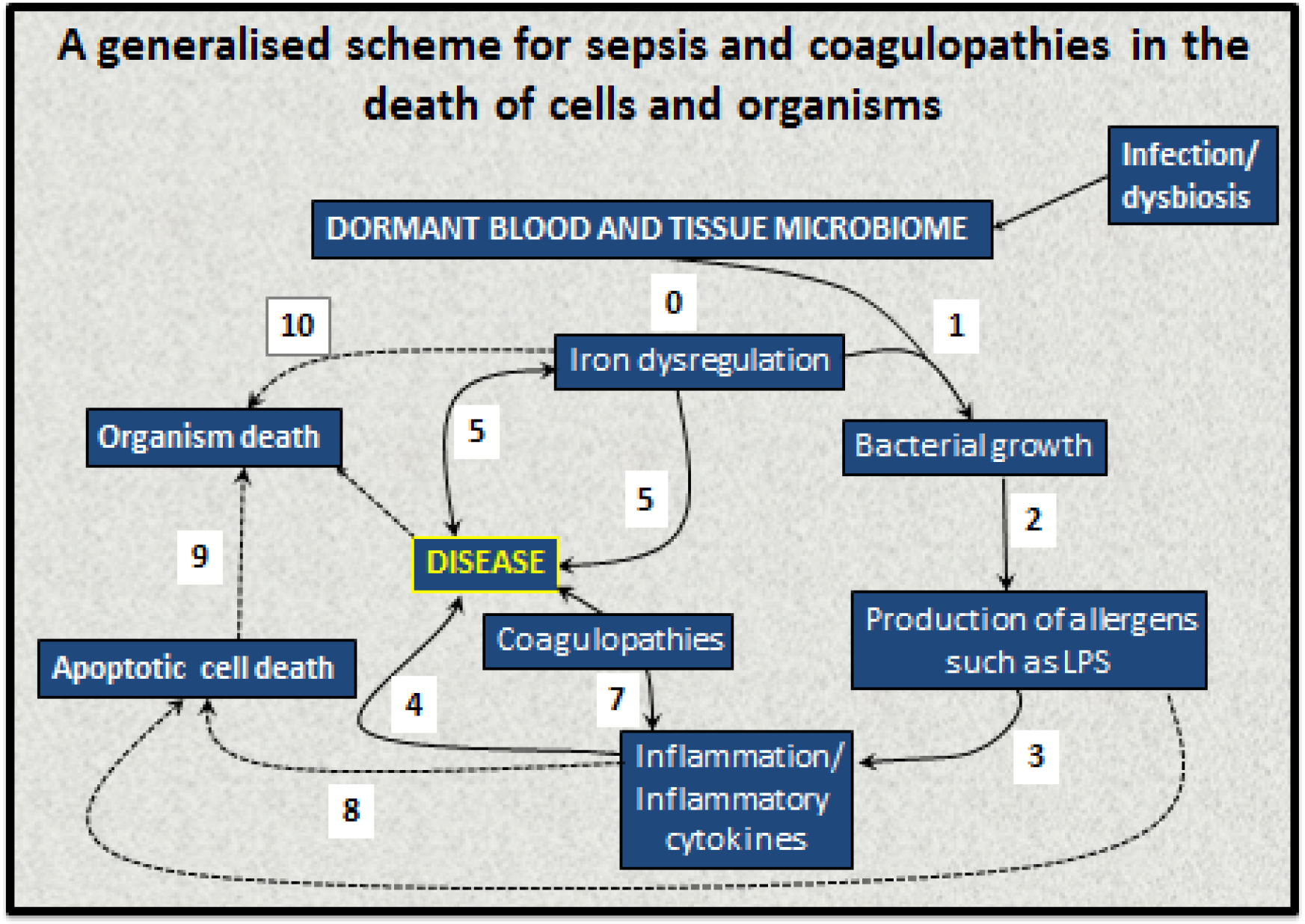
A systems biology model of the development of coagulopathies during SEPSIS, SIRS and MODS.

Recombinant soluble human thrombomodulin (TM-α) is a novel anticoagulant drug, and has been found to have significant efficacy in the treatment of sepsis-based DIC [192–201], again adding further weight to our hypothesis. As Okamoto and colleagues [178] point out, “In the European Union and the USA, the 2012 guidelines of the Surviving Sepsis Campaign do not recommend treatment for septic DIC [3; 202]. In contrast, in Japan, aggressive treatment of septic DIC is encouraged [203–205]”, and that “that Japan is one of the countries that most effectively treats patients with septic DIC” [178].

Thus, if it is accepted that the type of fibrin that is formed is substantially of the β-amyloid variety, then anticoagulant and other drugs that inhibit or reverse such amyloid processes should also be of value [206], as they seem to be in Alzheimer-type dementia [207–209].

## Concluding remarks

There is by now abundant evidence that coagulopathies involving fibrin clots are a major part of sepsis, SIRS, septic shock, MODS, DIC and organismal death. We have invoked further evidence that the type of fibrin involved is a β-amyloid form, and that it is this that is especially damaging; this definitely needs to be tested further, for instance using appropriate stains [127] and/or X-ray measurements [210] in concert with cellular toxicity assays. Finally, we suggest that anticoagulant therapies that inhibit or reverse those amyloid forms of fibrin production will be especially valuable. To this end, lowering the levels of fibrinogen itself would seem to be a desirable aim [211].

## Acknowledgments

We thank the Biotechnology and Biological Sciences Research Council (grant BB/L025752/1) as well as the National Research Foundation (NRF) and Medical Research Council (MRC) of South Africa for supporting this collaboration. This is also a contribution from the Manchester Centre for Synthetic Biology of Fine and Speciality Chemicals (SYNBIOCHEM) (BBSRC grant BB/M017702/1). We thanks Dr Janette Bester for confocal microscopy. DBK thanks Prof Nigel Harper for a useful discussion.

## Ethical approval disclosure

Ethical approval was granted at the University of Pretoria for all human studies (Human Ethics Committee: Faculty of Health Sciences): E Pretorius.

